# A glycosaminoglycan degrading Polysaccharide Utilization Locus in the oral microbiota species *Segatella oris*

**DOI:** 10.64898/2026.10.07.757250

**Authors:** Abel Cousin, Yara Awad, Rebecca-Joe Boustany, Rosa Lorizolla Cordeiro, Mélanie Friedel-Arboleas, Sophie Drouillard, Mélanie Loiodice, Laurine Buon, Chrystel Lopin-Bon, Rebekka Wild, William Helbert, Romain R Vivès, Marie Couturier

**Author notes:** Equivalent contribution (Abel Cousin, Yara Awad).

## Abstract

The human gut microbiota has been explored for the last two decades and studies have uncovered its major role in the degradation of complex glycans from the diet as well as the host tissues. Strikingly, until very recently, no similar investigation has been carried out on the human oral cavity microbiota, which is increasingly recognized has an important parameter in gastrointestinal health. *Prevotellaceae* are an important and under-investigated family of bacteria in both the gut and the oral cavity microbiota. The oral cavity is particularly rich in glycosaminoglycans which could provide an abundant carbon source for the inhabiting microorganisms, and play a significant role in host-pathogen interactions. Hence, we analyzed Polysaccharide Utilization Loci in oral microbiota species and we identified a glycosaminoglycan-degrading Polysaccharide Utilization Locus in *Segatella oris*. We produced and characterized the four enzymes, confirmed their ability to degrade glycosaminoglycans, and we focused our efforts on the biochemical and structural characterization of the first reported *Prevotellaceae* endosulfatase.

## Introduction

The gastrointestinal tract (GIT) microbiota is composed of complex populations of microorganisms inhabiting the human body between the oral cavity and the colon ^1^. In the GIT, the *Bacteroidales* order is a major taxonomic group in terms of abundance and effects exerted on the human body ^2^. In this order of bacteria, two closely related families, *Bacteroidaceae* and *Prevotellaceae*, are alternatively dominating the bacterial populations ^3^. For instance, *Bacteroidaceae* are dominant in the gut in individuals with a westernized diet, whereas *Prevotellaceae* such as *Segatella copri* dominate high-fiber rich and vegetarian diet microbiota, generally recognized as healthy gut microbiotas ^4,5^. However, in the oral cavity, bacterial populations differ, since *Prevotellaceae* has been described as the most abundant family among *Bacteroidales*, especially in saliva, both in healthy and dysbiosed oral microbiotas ^6,7,8^.

Over the last two decades, the functional characterization of the gut microbiota and its impact on many aspects of human health has been massively investigated. In particular, the microbiota collectively degrades complex polysaccharides from the diet and the host tissues, providing numerous benefits to the host such as energy acquisition ^9^, production of vitamins ^10^ and short chain fatty acids ^11^, development and modulation of the immune system maturation ^12^. Host-derived polysaccharides, such as glycosaminoglycans (GAGs), are a constant and stable source of carbon abundantly present in mammalian tissues and are likely degraded throughout the gastrointestinal tract (GIT) ^13^. In the gut, *Bacteroiaceae* are the main degraders of GAGs and mucins ^14,15^ whereas *Segatella copri*, which is the most abundant *Prevotellaceae* in the gut, is not able to forage these glycans and is rather specialized in cereal fibers utilization ^16^.

Contrary to the gut microbiota, the functional role of the oral microbiota and its capacity to degrade glycans has been largely underinvestigated so far. Recently, the role of this specific community in oral and systemic health has emerged. In particular, as bacteria can be easily translocated from the mouth to the respiratory and digestive tracts, it is now considered that the oral microbiota composition might exert an effect on health, both locally and at distant sites ^17,8^. In particular, it has been recently suggested that some oral bacteria are able to use GAGs for their growth. For instance *Granulicatella adiacens*, a species found in oral microbial communities, was shown to grow on hyaluronic acid ^13^. To the best of our knowledge, no investigation has been performed at the molecular level to evaluate the degradation of GAGs by the oral microbiota bacteria, of which *Prevotellaceae* are important members.

GAGs are composed of repeated disaccharide units comprising a uronic acid or galactose residue and an amino sugar. Variations in the type of uronic acid (glucuronic acid or iduronic acid), amino sugar (*N*-acetylglucosamine or *N*-acetylgalactosamine) and position and number or sulfate decorations define several classes of GAGs (heparin/heparan sulfate, chondroitin sulfate/dermatan sulfate, keratan sulfate, and hyaluronic acid) ^18^. GAG degradation by microbiota species requires several enzymes with complementary activities, including Polysaccharide Lyases (PL), Glycoside Hydrolases (GH) and sulfatases. PL degrade the GAG chains and release disaccharides, which are further processed into monosaccharides by GH. Both PL and GH belong to the Carbohydrate Active enZymes (CAZymes) and as such, are classified in the CAZy database (www.cazy.org, ^19^). Sulfatases, which catalyze the removal of sulfate groups from glycan chains, are classified in the SulfAtlas database (https://sulfatlas.sb-roscoff.fr, ^20^).

In the *Bacteroidetes* phylum, the genes encoding CAZymes that target a specific glycan are colocalized and co-regulated in Polysaccharide Utilization Loci (PULs), together with proteins involved in glycan sensing, capture and oligosaccharide internalization. PULs are classified in the PULDB extension of the CAZy database (www.CAZy.org/puldb, ^21^. PULs from gut *Bacteroidaceae* have been extensively studied, which helped to uncover the ability of several species to degrade multiple complex glycans from the diet or from the host tissues ^22,23,24^. In particular, GAG enzymatic degradation has been extensively dissected in the *Bacteroidaceae* model bacterium *B. thetaiotaomicron* ^25,15^. On the contrary, the glycan degrading ability of the iconic *Prevotellaceae* species *Segatella copri* have just started to be investigated at the molecular level, with several dietary plant polysaccharide degradation systems being elucidated ^26,27,28^. To date, no study has detailed the role of host glycan targeting PULs in oral *Prevotellaceae*.

Here, we analyzed putative GAG-targeting PULs in *Prevotellaceae* and we identified one in *Segatella oris* (formerly *Prevotella oris* ^29^), a member of the oral microbiota. Production and purification of the three CAZymes and the sulfatase of the PUL was achieved, and their activity as well as their complementary actions in GAG degradation could be assigned. As sulfatases are key enzymes in the degradation of GAGs, we particularly focused our efforts on the biochemical and structural characterization of *S. oris* endo sulfatase *So*S1_27, the first characterized *Prevotellaceae* endosulfatase.

## Results

### In *Prevotellaceae*, GAG-degrading PULs are restricted to species from the oral cavity

The PULDB gathers thousands of predicted PULs from the *Prevotellaceae and Bacteroidaceae* families. A search for PULs containing at least (i) one sulfatase and (ii) one gene from a known GAG-degrading PL family (PL8, PL12, PL15, PL29, PL33, PL35) showed that the ability to degrade GAGs seems to be more common among *Bacteroidaceae* than *Prevotellaceae* (Supplementary Table S1). *Segatella copri* is the emblematic *Prevotellaceae* species, most frequently identified in the gut. Strikingly, no GAG-degrading PUL could be identified in *S. copri*. Analysis of *S. copri* annotated CAZomes (i.e. ensemble of predicted CAZymes), that are available publicly in the CAZy database, showed that some strains of *S. copri* do encode one single PL33 (www.cazy.org, ^19^). This family of lyases has recently been subjected to in-depth characterization ^30^ and revealed a variety of specificities for GAGs, suggesting that some degradation of GAGs by *S. copri* might be possible. However, PUL_GAG_ in *Prevotellaceae* seem to be restricted to species that are primarily from the human oral microbiota, e.g. *S. oris, P. maculosa*, *P. dentalis* (Figure 1). Based on this analysis, we selected *Segatella oris* GAG-degrading PUL (PUL_GAG_) for biochemical characterization.

**Figure 1:**
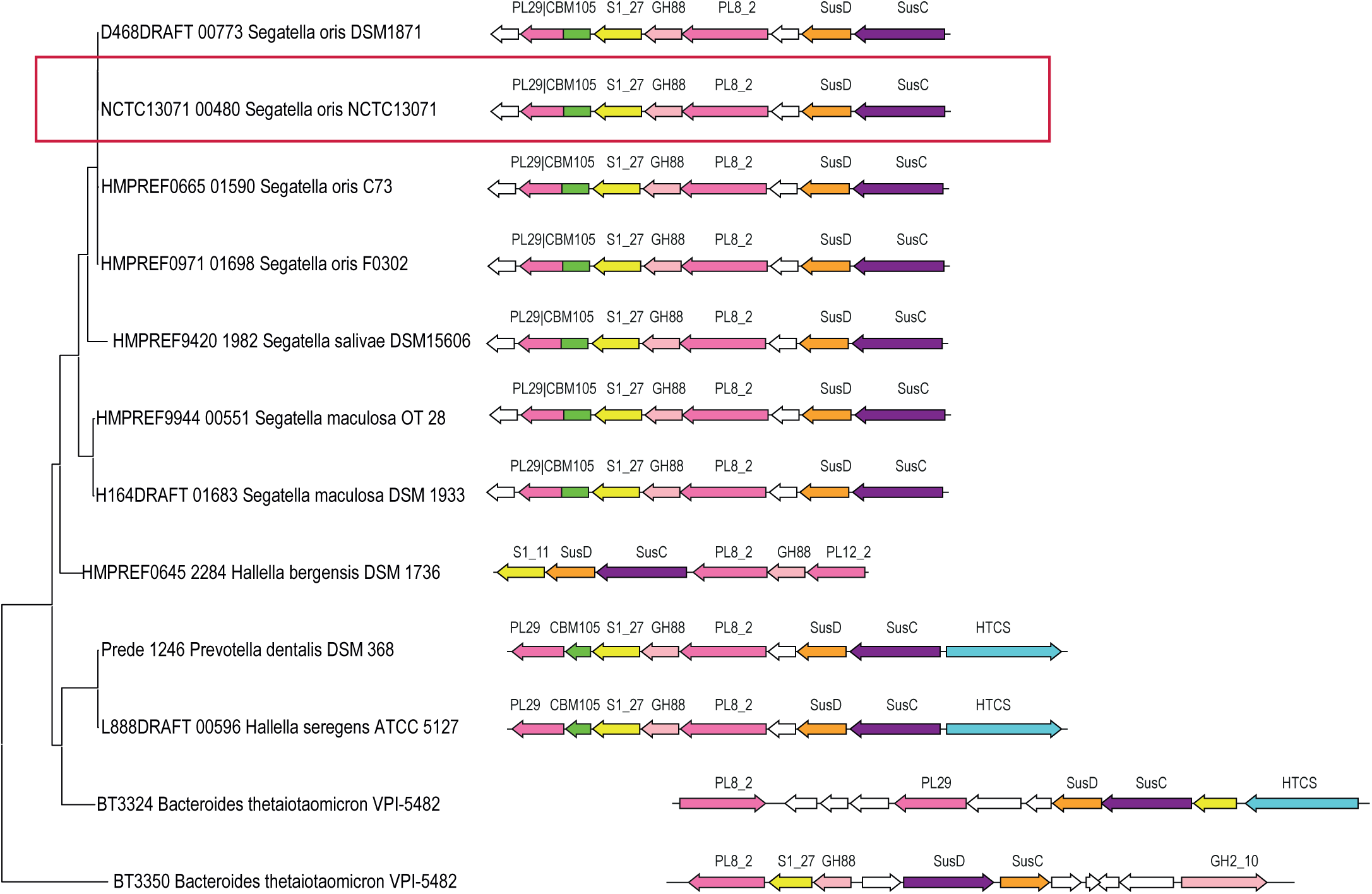
PUL_GAG_ organization in *Prevotellaceae* species. *Prevotellaceae* PUL_GAG_ were extracted from PULDB and a distance tree was build based on PL8 sequences. The selected *S. oris* PUL is highlighted in a red frame.

### Organization of *S. oris* GAG degradation PUL

*S. oris* PUL_GAG_ comprises eight genes, of which three are predicted to encode CAZymes from family PL8, PL29 and GH88, respectively (Table 1, Figure 1). One gene is predicted to encode a carbohydrate sulfatase belonging to subfamily S1_27 of the SulfAtlas classification (https://sulfatlas.sb-roscoff.fr ^20^. The PUL also encodes a putative TonB-dependent transporter (SusC homolog) and a putative surface glycan-binding protein (SusD homolog), both proteins being considered a hallmark for the presence of PULs. Two proteins of unknown function are also encoded by *S. oris* PUL_GAG_. BLAST analysis indicates that these sequences contain predicted domains of unknown function (DUF4861 and DUF5017, respectively). No DUF4861 containing protein was found in the gut model bacterium *B. thetaiotaomicron*, whereas a DUF5017 containing protein is also encoded in *B. thetaiotaomicron* PUL_GAG,_ where it was identified as a SGBP glycan binding protein ^15^. Overall, comparison with the PUL_GAG_ from *B. thetaiotaomicron*, which expresses two PL8 enzymes with complementary modes of action ^15^, as well as two carbohydrate sulfatases, highlights*S. oris* PUL as a condensed version of the GAG-degrading locus found in *Bacteroides* species. In *B. thetaiotaomicron*, a PL33 gene that is located outside of the PUL has been shown to be co-expressed with the PUL and to be active on GAGs. In the *S. oris* NCTC13071 genome, a PL33 gene is also encoded outside the PUL. In an earlier work, we characterized the corresponding enzyme that showed low activity on hyaluronic acid (HA), chondroitin sulfate (CS) and dermatan sulfate (DS) ^30^.

**Table 1:** Characteristics of *S. oris* PUL_GAG_ encoded proteins.

| GenBank | Protein | Predicted family | Predicted enzymatic activity | Signal peptide * | *B. thetaiotaomicron* homologous proteins (% identity) |
| --- | --- | --- | --- | --- | --- |
| VEH14499.1 | DUF4861 domain-containing protein | na | na | No | No detected similarity |
| VEH14500.1 | *So*PL29 | PL29-CBM105 | GAG lyase | Yes | AAO78434.1 (41%) |
| VEH14501.1 | *So*S1_27 | Sulf1_27 | Carbohydrate sulfatase | Yes | AAO78455.1 (66%) |
| VEH14502.1 | *So*GH88 | GH88 | Unsaturated glucuronyl hydrolase | Yes | AAO78454.1 (63%) |
| VEH14503.1 | *So*PL8 | PL8_2 | GAG lyase | Yes | AAO73430.1 (57%) |
| VEH14504.1 | DUF5017 domain-containing protein | na | na | Yes | AAO78436.1 (48%) |
| VEH14505.1 | SusD | na | na | No | AAO78437.1 (61%) |
| VEH14506.1 | SusC | na | na | Yes | AAO78438.1 (67%) |

na : not applicable. \*Signal peptide prediction using Phobius (https://phobius.sbc.su.se/)<sup>57</sup>

### Production and activity screening of *S. oris* PUL_GAG_ enzymes

The four predicted enzymes encoded by *S. oris* PUL_GAG_ were successfully produced in *E. coli* and purified by one or two steps of chromatography. To evaluate their substrate preference, the four enzymes were incubated with a library of GAG polysaccharides and oligosaccharides and degradation products were analyzed using size-exclusion chromatography (Table 2 and Figure 2, see also Supplementary Table S2 for a list of substrates used in the study). This activity screening confirmed that *S. oris* PUL_GAG_ enzymes target GAGs. The two putative PL were found to degrade HA, CS and DS, releasing oligosaccharides upon incubation with the polysaccharide substrates (Figure 2A). GH88 enzymes are unsaturated glucuronyl hydrolases which further degrade unsaturated oligosaccharides released by PL. As expected, *So*GH88 was active on all non-sulfated or desulfated GAG oligosaccharides that were tested (Figure 2B). The fourth enzyme of the PUL, the putative sulfatase *So*S1_27, was first assayed on the synthetic substrate paranitrophenol-sulfate to verify the proper maturation of the enzyme in *E. coli*, which indeed confirmed that *So*S1_27 was properly folded and maturated (Table 2). Further screening showed that *So*S1_27 was able to release sulfate from both CS and DS poly– and oligosaccharides, as a peak compatible with a sulfate group was observed upon incubation of these substrates with the enzyme (Figure 2A, 2B).

**Figure 2:**
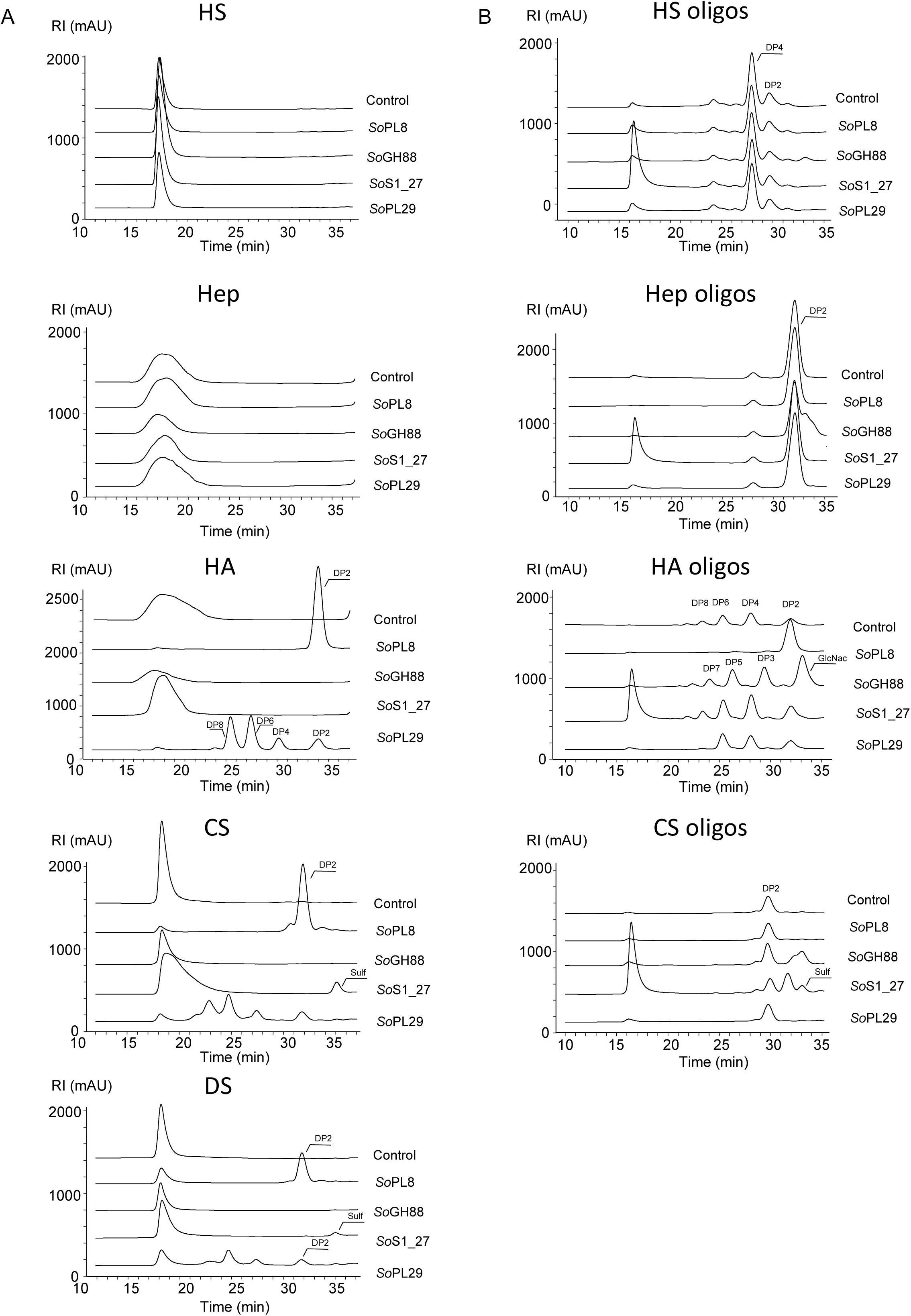
Activity of *S. oris* PUL_GAG_ enzymes on GAG oligo– and polysaccharides. Each enzyme of the PUL was tested on a range of GAG substrates. Panel A: polysaccharides, Panel B: oligosaccharides. HS: heparin sulfate, Hep: heparosan, HA: hyaluronic acid, CS: chondroitin sulfate, DS: dermatan sulfate.

**Table 2:** Activity of *S. oris* PUL_GAG_ enzymes on various GAG substrates.

| Enzyme | Synthetic substrates | oligosaccharides | Polysaccharides |
| --- | --- | --- | --- |
| *So*PL8 | na | CS, DS | CS, DS, HA |
| *So*S1_27 | pNP-S | CS, DS | CS, DS |
| *So*PL29 | na | nd | CS, DS, HA |
| *So*GH88 | na | CS, DS, HA, Hep, HS | nd |

na: not assayed. nd: not detected

### Analysis of *So*PL8 and *So*PL29 mode of action and sequential activity of the four enzymes

Next, we investigated the mode of action of the two lyases. Exo-acting lyases release only disaccharides from the extremities of the polysaccharide chain, whereas endo-acting enzymes randomly cleave throughout the glycan chain, first releasing large oligosaccharides which size decreases over the course of the reaction. To determine the endo or exo mode of action of the two lyases *So*PL8 and *So*PL29, degradation of HA was monitored over time, and digestion products were analyzed using size exclusion chromatography (Figure 3). This showed that *So*PL29 is an endo-active PL, first releasing large oligosaccharides whose size decreases over the course of the reaction (Figure 3A), whereas *So*PL8 showed a complementary exo mode of action, producing only DP2 from the extremity of the chains (Figure 3B).

**Figure 3:**
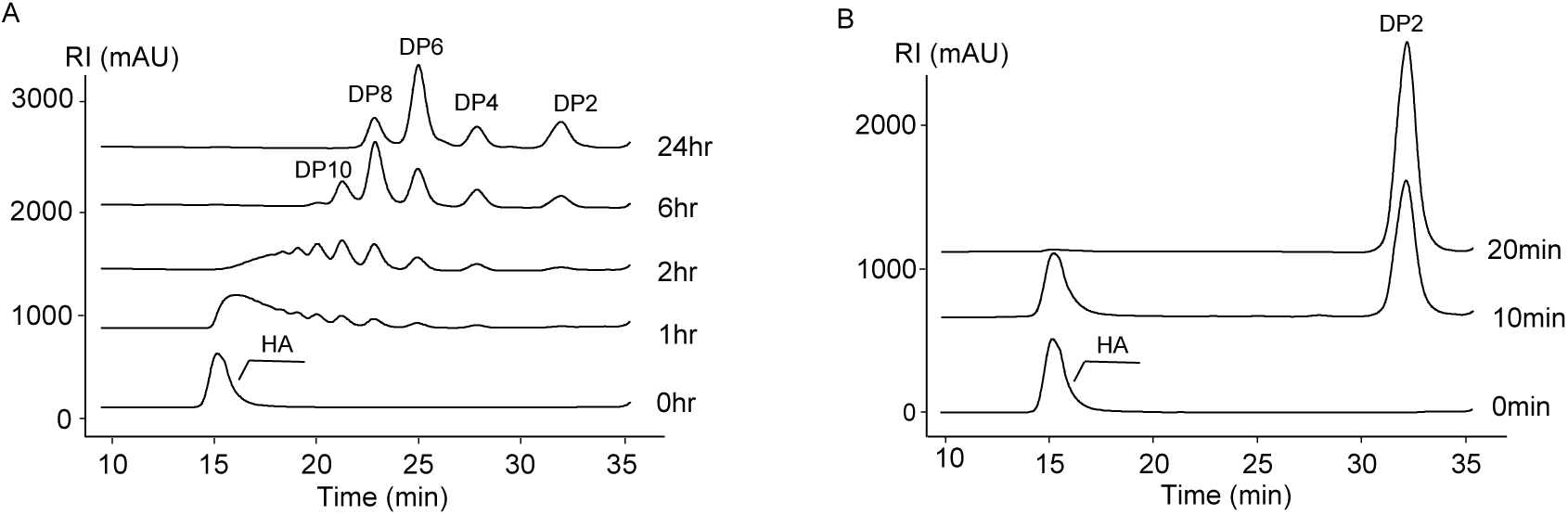
Mode of actions of *So*PL29 and *So*PL8 on hyaluronic acid. A: HPLC-SEC product profiles generated over time by the degradation of hyaluronic acid by *So*PL29. B: HPLC-SEC product profiles generated over time by the degradation of hyaluronic acid by *So*PL8.

Enzymes encoded by PULs usually degrade a glycan in a cooperative manner. The screening results showed that the four enzymes encoded by *S. oris* PUL target HA and CS or their degradation products. Hence, we next assessed the sequential activity of the four enzymes on CS to confirm the cooperative degradation pathway. SEC analysis showed that CS was efficiently degraded by the sequential action of the four enzymes to a mixture of monosaccharides (Figure 4).

**Figure 4:**
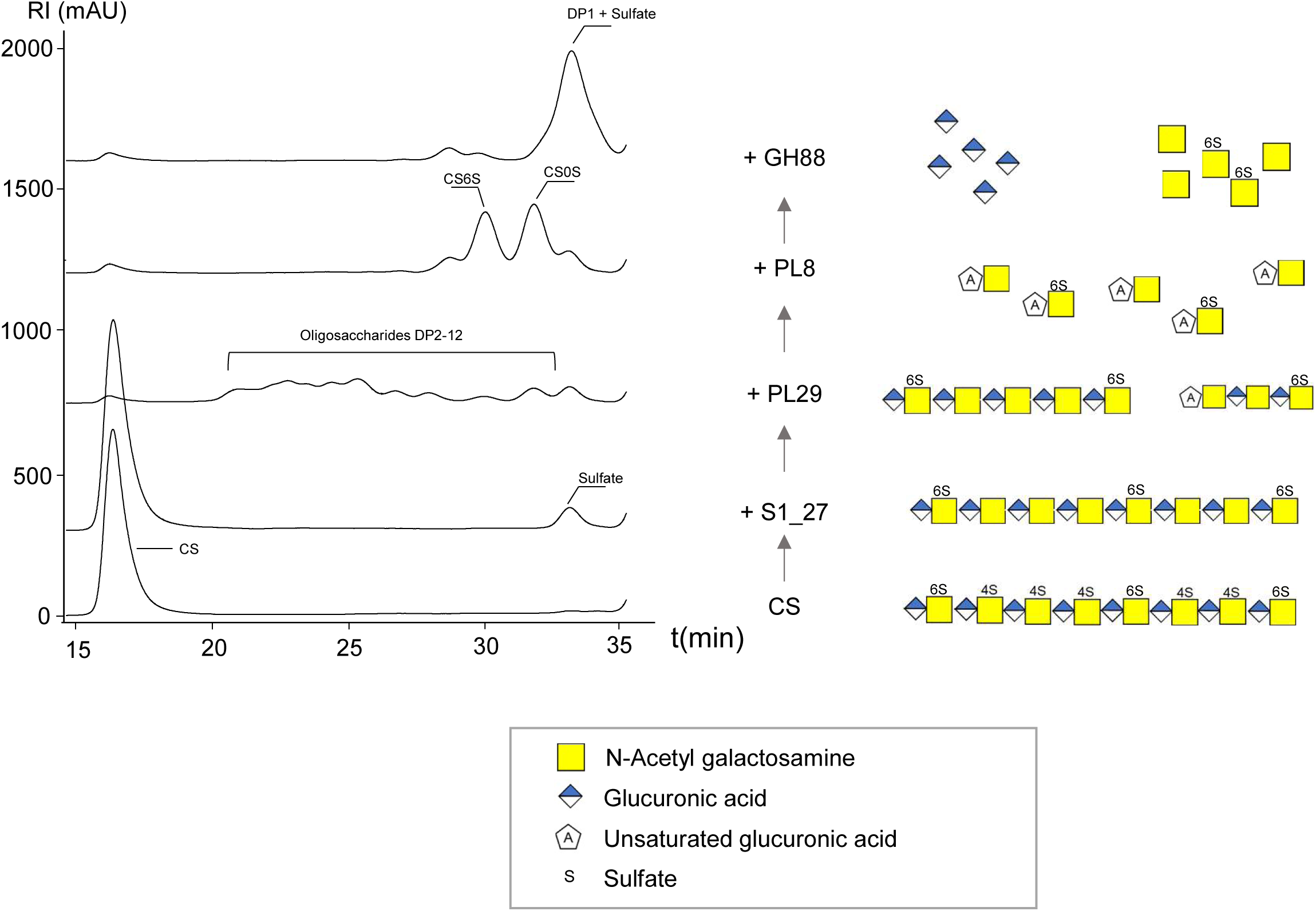
Sequential degradation of CS by the four enzymes of *S. oris* PUL. HPLC-SEC product profiles generated upon sequential degradation of CS by the four enzymes of the PUL. Samples were taken after 24 hours incubation with each enzyme and before addition of the next enzyme.

### Biochemical characterization of *So*S1_27

Since the activity screening had suggested that *So*S1_27 was active on CS poly– and oligosaccharides, we then aimed at assessing whether *SoS1_27* was an endo– or exo-active enzyme. We first compared the profile of oligosaccharides produced by *So*PL29 on *So*S1_27-pretreated CS versus untreated CS (Supplementary Figure S1). The differences in the resulting SEC profiles suggested extensive desulfation consistent with an endo-active enzyme, since all oligosaccharide sizes were shifted upon *So*S1_27 desulfation, compared to the untreated control. H^1^ NMR analysis of desulfated CS was then performed, showing a complete loss of the 4S signal and therefore confirming the endo activity of *So*S1_27 (Supplementary Figure S1). The biochemical characterization of *So*S1_27 on a range of chondroitin disaccharides with defined sulfation patterns (CS4S, CS4S6S, CS2S6S, CS2S4S6S, see Supplementary Table S2) confirmed that the enzyme specifically removes 4-O sulfation from both monosulfated CS4S and disulfated CS4S6S disaccharides, (Figure 5, Table 3), whereas no activity was observed on CS2S4S, CS2S6S or CS2S4S6S disaccharides.

**Figure 5:**
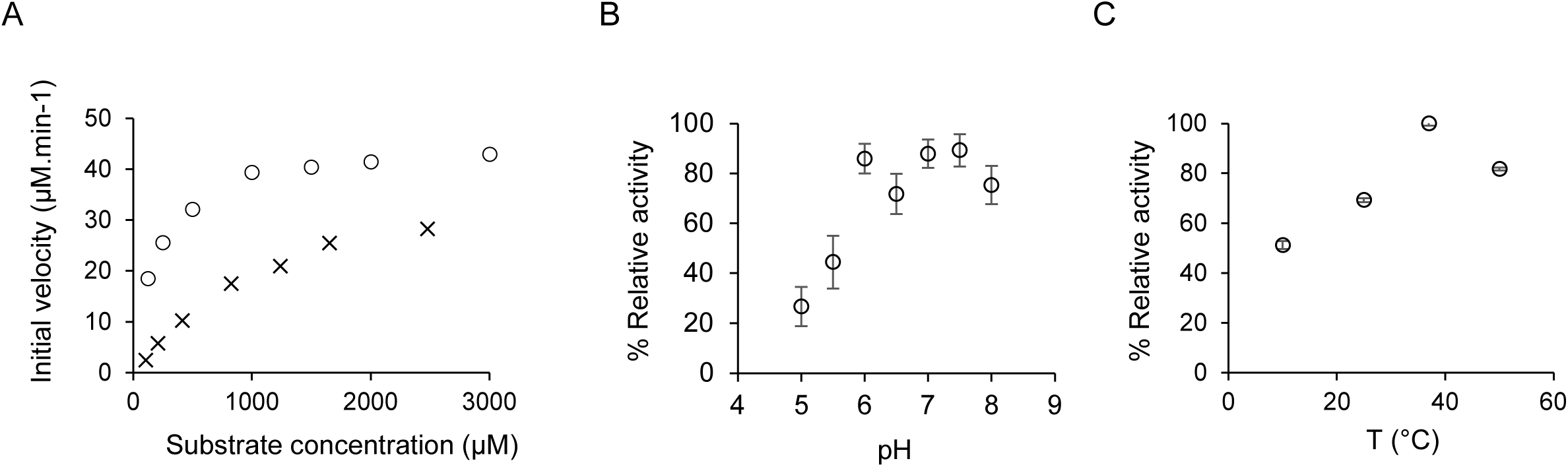
Biochemical characterization of *So*S1_27. A: Initial velocities of disaccharide desulfation as a function of substrate concentration. o: CS4S, x: CS4S6S. B: optimal pH was determined on CS4S between pH5 and 8, C: Optimal temperature was determined on CS4S between 10 and 50°C.

**Table 3:** Kinetic parameters of *So*S1_27 on chondroitin sulfate oligosaccharides.

| Disaccharide | Activity | V<sub>max</sub>(μM.min-1) | K<sub>cat</sub>(s-1) | K<sub>M</sub>(μM) | K<sub>cat</sub>/K<sub>M</sub>(s-1.μM-1) |
| --- | --- | --- | --- | --- | --- |
| CS4S | Yes | 34.4 +/- 14.5 | 11.5 +/- 4.8 | 211 +/- 40.3 | 5.4 x 10<sup>-2</sup> +/- 2.6 x 10<sup>-2</sup> |
| CS4S6S | Yes | 46.5 +/- 12.4 | 3.9 +/- 1.0 | 1380 +/- 244 | 2.8 x 10<sup>-3</sup> +/- 9.2 x 10<sup>-4</sup> |
| CS2S4S | No |  |  |  |  |
| CS2S4S6S | No |  |  |  |  |

Initial velocities of recombinant *So*S1_27 were determined on CS disaccharides CS4S and CS4S6S using increasing concentrations of substrates, enabling determination of kinetic parameters (Figure 5A, Table 3). The catalytic efficiency was an order of magnitude higher for CS4S than CS4S6S, with 5.4 x 10-2 µM^-1^.s^-1^ for CS4S versus 2.8 x 10-3 µM^-1^.s^-1^ for CS4S6S. We also investigated the optimal temperature and pH for *So*S1_27 sulfatase activity on its preferred substrate CS4S, and measured an optimal activity at 37°C with more than 70% activity retained between 25 and 50°C, and an optimal pH around 7, with 60% activity observed between pH 6 and 8 (Figure 5, B-C).

### Crystal structure of *So*S1_27

To learn more about the architecture of *So*S1_27 and the spatial organization of its catalytic site, we determined its structure at 2.4 Å resolution (space group P1, R-work = 0.2141, and R-free = 0.2746) using X-ray crystallography (Supplementary Table S3). We found four molecules in the asymmetric unit, with pairwise RMSD values ranging from 0.217 to 0.62 Å across 451 Cα atoms (Supplementary Figure 2A). PISA analysis ^31^ suggests that the crystallographic arrangement is unstable, which was also supported by mass photometry experiments indicating that *So*S1_27 is predominantly monomeric in solution (Supplementary Figure 2B). The structure reveals that *So*S1_27 adopts a fold typically found in S1 sulfatases with a two-domain architecture ^32^. The N-terminal domain contains a central parallel β-sheet, with β-strand 7 facing the other direction, and surrounded by α-helices. The C-terminal domain has a four stranded anti-parallel β-sheet, flanked by one longer helix (Figure 6A).

**Figure 6.**
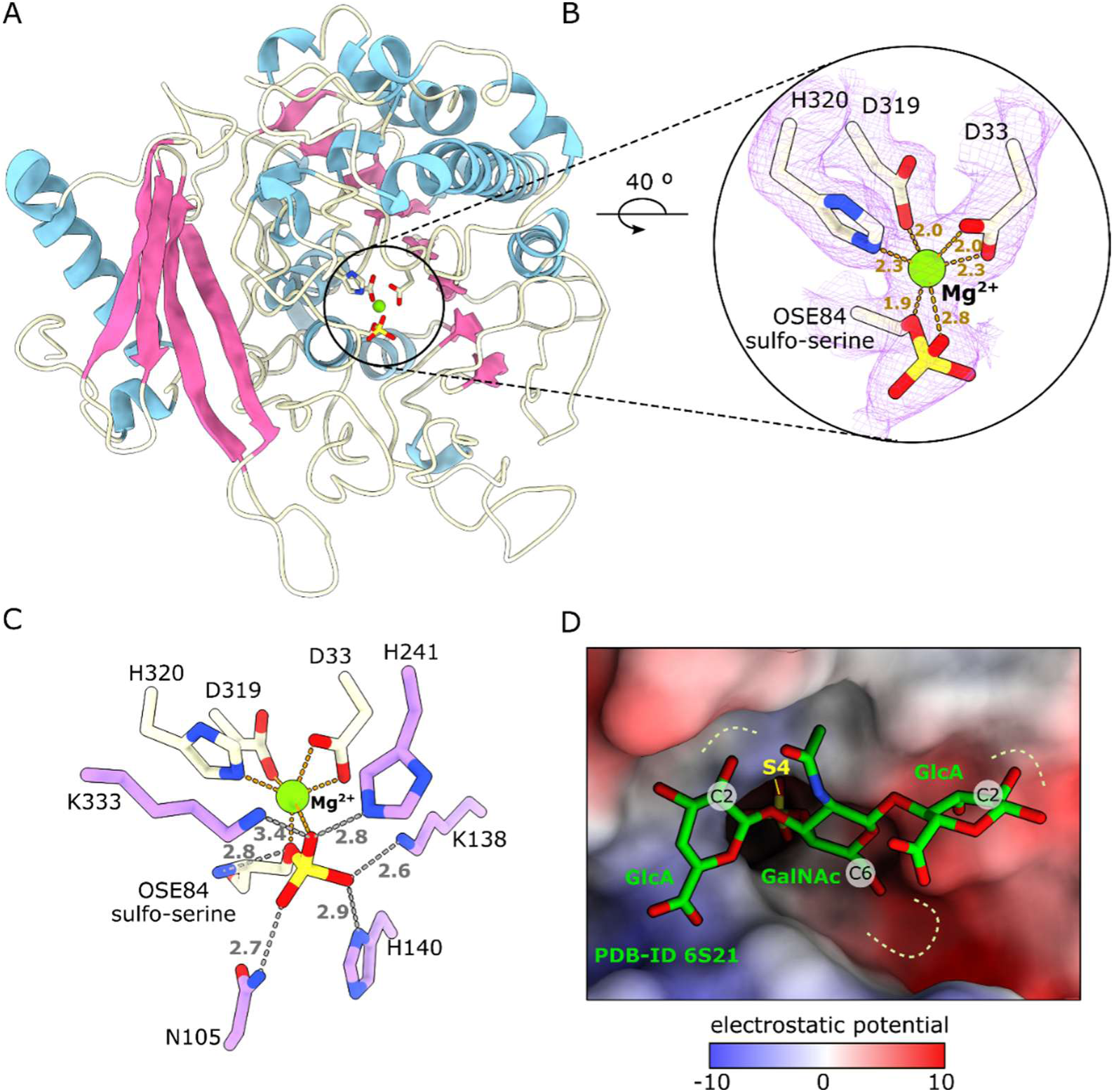
Crystal structure of *So*S1_27. A: Overall folding represented in secondary structure, in which the α-helices are colored in light blue, the β-sheets in pink, and the loops in light yellow. B: A detail of the metal ion-binding site, with the side chains of the residues coordinating Mg2+ (green sphere) represented in sticks. The 2Fo-Fc map, contoured at 1.5 σ, is shown as a purple mesh. C: Zoom-in on important interactions in the active site, harboring the sulfate, which is covalently-bound to the catalytic serine (OSE84). The residues interacting with the sulfate group are shown as purple sticks and residues interacting with the metal-ion are shown as light yellow sticks. The Mg2+ ion is shown as green sphere. D: Superposition of the *So*S1_27 structure (surface, colored by electrostatic potential) and the CS-derived trisaccharide bound to BT3349 from *B. thetaiotaomicron* (sticks, green carbons) (PDB-ID 6S21). The sulfo-serine OSE84 was removed from the *So*S1_27 structure for a better illustration of the sulfate binding pocket.

We identified the active site based on homology with other FGly-dependent sulfatases. Notably, the residues that are involved in metal ion coordination (Asp33, Asp319 and His320) (Figure 6B) are conserved within the S1_27 subfamily (Supplementary Figure 2C). We interpreted the metal ion as Mg^2+^, despite the fact that calcium is commonly observed in S1 sulfatases. Importantly, Mg^2+^ was present in the crystallization solution, whereas calcium ions were not supplied during protein expression or purification. The identity of the Mg^2+^ was further supported by analysis using the CheckMyMetal server ^33^.

Despite soaking of the crystal with a CS-derived tetrasaccharide, a substrate analog of *So*S1_27, no clear electron density for oligosaccharides was detected in the structure. Instead, a continuous electron density was observed around the side chain of the catalytic serine (Ser84), consistent with a covalently-linked sulfate group (Figure 6B-C). Such a covalent sulfo-serine reaction intermediate has been described for various S1 sulfatases ^34^. In the *So*S1_27 structure, it forms interactions with the side chain of five amino-acid residues, the main chain of Sulfo-Ser84 (OSE84) and the coordinated metal ion in the active site (Figure 6C). We then compared the *So*S1_27 structure to that of the endosulfatase BT3349 from *B. thetaiotaomicron*. The two sequences share 66% sequence identity, and the structure of *B. thetaiotaomicron* was solved bound to a CS-derived trisaccharide containing two GlcA and one GalNAc units, and a sulfate group at the C4 position of GalNAc (PDB-ID 6S21 ^15^) (Figure 6D). The comparison of our structure with that of BT3349 showed that *So*S1_27 conserves a small pocket for accommodating and cleaving the sulfate group at C4 position in GalNAc residues.

## Discussion

The human microbiota plays a central role in health and disease, yet the contribution of oral microorganisms remains comparatively underexplored. Indeed, while it is well established that the gastrointestinal tract (GIT) microbiota is a key parameter of human health, the oral microbiota has only recently been recognized as an important player ^8,9^. In particular, the oral-gut axis is now well established, underscoring the influence of this microbial community on distal body sites ^17^. Taxonomic studies have highlighted that *Prevotellaceae* are identified in both healthy and dysbiotic oral cavity microbiota, where their relative abundance can reach 10 to 15% of total bacteria ^35^. Recent research also highlighted cross-feeding interaction between *S. oris* and the pathogenic bacteria *Fusobacterium nucleatum* ^36^. Interestingly, the oral cavity is particularly rich in mucins and GAGs, suggesting that these glycans may represent a key carbon source for inhabiting bacteria, but functional studies are missing.

Genomic analyses have established that *Prevotellaceae* encode a large number of polysaccharide utilization loci (PULs). However, only a few of these PULs are predicted to target host-derived glycans, suggesting that the repertoire of glycan-degrading enzymes differs substantially from that of canonical gut commensals. In particular, we observed that GAG-targeting PULs seem to be restricted to oral species of the *Prevotella* and *Segatella* genera, and are not found in the iconic gut-enriched *S. copri* species. This observation is consistent with previous reports of host-specific adaptations within this genus where distinct PUL and CAZyme repertoires reflect differential glycan utilization strategies ^35,37^.

Our characterization of the PUL_GAG_ from *Segatella oris* confirmed that this oral species encodes a set of enzymes able to efficiently break down HA, CS and DS. This PUL includes in particular two PL (PL29 and PL8) and one GH (GH88) with complementary activities. During the course of this work, *So*PL29 was studied by another group, showing a modular structure containing a PL29 domain attached to a Carbohydrate Binding module (CBM) belonging to a new family and termed CBM105. In this work, the authors described *So*PL29 substrate specificities, and showed that the enzyme tackles efficiently HA and CS, with a random endo-acting mode, and these conclusions were in agreement with our observations^38^. The PL33 gene located outside of the PUL could work synergistically with the other lyases, as biochemical characterization showed that the preferred substrate of this enzyme was HA ^30^. The gut model bacterium *B. thetaiotaomicron* GAG degrading PUL, which has been extensively characterized, also encodes a PL33 that takes part in the degradation of GAGs ^15^. *Bacteroides* and *Segatella* are closely related genera and it is not surprising that they share related metabolic capabilities. It is however interesting to note that the ability to degrade GAGs throughout the gastrointestinal tract is not dominated by one single taxonomic group, but is rather distributed between *Segatella* in the oral cavity and *Bacteroides* in the gut.

The identification of *So*S1_27 as the first biochemically and structurally characterized sulfatase from *Prevotellaceae* GIT microorganism highlights the need for deeper investigation of *Prevotellaceae* enzymes to clarify their roles in the GIT microbiota. Sulfatases are critical for GAG metabolism in both eukaryotic and prokaryotic systems. Endosulfatases are particularly noteworthy, as they catalyze internal polysaccharide modifications rather than stepwise exolytic removal. In mammals, the extracellular sulfatase Sulf2 modulates the structure of heparan sulfate, exerting profound effects on signaling pathways ^39^. In bacteria, endosulfatase activity contributes to efficient GAG assimilation and community dynamics within the gut ^15^. In this context, S*o*S1_27 represents a unique link between oral bacterial metabolism and host glycan remodeling, with potential consequences for microbial competition and host–microbiota interactions. Similar endosulfatase activity has been observed in gut-associated bacterial sulfatases ^40^, and subsequent structural studies have confirmed the active site architecture ^15^, highlighting the conservation of this mechanism across microbiota-encoded enzymes.

Analysis of *So*S1_27 three-dimensional structure and comparison with that of *B. thetaiotaomicron* in complex with a CS-derived trisaccharide showed that *So*S1_27 harbors a pocket accommodating the sulfate group at the C4 position, thus allowing sulfate hydrolysis. On the contrary, the interaction surface between C2 carbon of GlcA and *So*S1_27 cannot accommodate a sulfate group, which would cause a steric hindrance unfavorable to the proper accommodation of the substrate, explaining the lack of activity on substrates containing sulfate groups at the C2 position. Furthermore, although the C6 carbon of the GalNAc portion fits into a small pocket that could accommodate a sulfate group, the negative surface charge suggests that this interaction would not be favorable. This may explain that although the enzyme can tolerate a sulfate group at the C6 position, there is one order of magnitude difference in the catalytic efficiency of *So*S1_27 towards CS4S6S as compared with CS4S. Overall, the identification and structural analysis of *So*S1_27 advance our understanding of oral *Prevotellaceae* biology and highlight endosulfatases as critical yet understudied mediators of host–microbiota interactions. By targeting internal sulfate groups within GAGs, *So*S1_27 may provide *S. oris* with a competitive advantage in the oral niche, influencing both microbial community composition and host mucosal ecology.

In general, our findings expand the repertoire of microbiota-associated enzymes involved in GAG metabolism. It is also, to the best of our knowledge, the first characterization of a glycan degrading system in the oral microbiota, which has recently emerged as a key community with great impact on health and disease. Future studies will be needed to address how these GAG degrading systems, and in particular sulfatases, impact oral or systemic health, and how these enzymes can be leveraged as biotechnological tools.

## Materials and Methods

### Bioinformatic analysis

A search for PULs in *Prevotellaceae* containing at least (i) one gene encoding a predicted carbohydrate sulfatase and (ii) one gene encoding a predicted GAG-targeting PL (PL8, PL12, PL15, PL29, PL33, PL35) was carried out in the PULDB^21^. Phylogenetic analyses of the set of PL8 sequences located in the PUL were conducted using MEGA version 12^41^. Sequences were aligned with MUSCLE ^42^, and a distance matrix was built from the multiple sequence alignment using the maximum likelihood.

### Gene synthesis

Sequences of putative CAZyme-encoding genes were retrieved using the PUL database (www.cazy.org/PULdb, [^21^]). The list of studied genes is given in Table 1. Gene sequences were optimized for expression in *Escherichia coli* and signal peptides if present were trimmed. Genes synthesis and cloning into the expression plasmid pET-28a (+), in frame with a His6-tag upstream of the coding sequence, were outsourced to Twist bioscience.

### Enzyme production and purification

Approximately 20 ng of each plasmid were used to transform E. coli BL21(DE3) competent cells. Clones harboring the recombinant expression plasmids were then grown in 5 ml LB precultures supplemented with 30 μg/mL kanamycin in a shaking incubator at 180 rpm and 37°C overnight. Cultures were subsequently carried out in 50 ml of NZY auto-induction LB media (NZYTech) medium supplemented with 30 μg/mL kanamycin, with 1/100 v/v overnight precultures and incubated at 37°C for 2 hours and then at 25°C for 22 hours. Cells were recovered by centrifugation at 4000g at 4°C for 5 min. To lyse the cells, the bacterial pellet was resuspended in buffer A (50 mm Tris pH7.8, 300 mM NaCl, 10 mM Imidazole) with 50 µg/ml lysozyme and stored at –80°C for one hour. After thawing, resuspended cells were further incubated with DNAse at 10°C for 15 min. Insoluble fractions were removed by centrifugation at 40,000g for 30 min at 4°C. The proteins were purified by affinity chromatography on a 1 mL HisTrap™ HP column (GE Healthcare) connected to a NGC chromatography system (Bio-Rad). The column was equilibrated with buffer A and the His6-tagged recombinant enzymes were eluted with an increasing gradient of imidazole between 0 and 300 mM. The fractions containing the proteins of interest were pooled and injected on a gel permeation ENRich650 column (Bio-Rad) and eluted in 20 mM Tris-HCl pH 7.5, 100 mM NaCl. The purity of the fractions was assessed using 10% SDS-PAGE analysis.

### Determination of enzymatic activity

#### Polysaccharide lyases

Enzyme activity was assayed on a range of glycosaminoglycans: Heparin sodium salt, heparosan, dermatan sulfate, hyaluronic acid and chondroitin sulfate sodium were tested following a protocol described earlier [^30,43^]. All substrates used for screening are presented in Supplementary Table S2. Briefly, 125µl of 0.4% substrate were incubated with appropriately diluted enzymes in Tris HCl 100 mM pH7.5, NaCl 50 mM with a final volume of 250µl. Enzyme concentrations ranged between 1 and 10 µM. Reactions were incubated overnight at 25°C and terminated by heating 10 minutes at 100°C, filtered through 0.2 µm membranes, and degradation products were analyzed by gel permeation chromatography using a Superdex peptide 10/300 (GE Healthcare) column connected to a high-performance liquid chromatography (HPLC) Ultimate 3000 system (Thermo Fisher). Injection volume was 50 µL and the elution was performed at 0.5 ml.min-1 in 0.1 M NaCl. Oligosaccharides were detected by differential refractometry (Iota 2 differential refractive index detector, Precision Instruments). When sequential reactions were performed, 0.2% substrate was incubated 24 hours successively with each enzyme in Tris HCl 100 mM pH7.5, NaCl 50 mM with a final volume of 1ml. 200 µl aliquots were taken at each time point to monitor substrate degradation. No enzyme activity was observed in heat-inactivated or enzyme-free control reactions, confirming that the observed activity was enzyme-dependent.

#### Unsaturated glucuronyl hydrolase

GH88 activity was assayed on GAG oligosaccharides produced in house using various GAG lyases. The enzymatic reaction contained 125µl of the produced GAG oligos with appropriately diluted enzymes in Tris HCl 100 mM pH7.5, NaCl 50 mM with a final volume of 250µl. The final enzyme concentration was 1-5 µM, and reactions were incubated overnight at 25°C with shaking. Analysis of the degradation product profiles was achieved by HPLC-SEC as described above. No enzyme activity was observed in heat-inactivated or enzyme-free control reactions, confirming that the observed activity was enzyme-dependent.

#### Sulfatase activity

Sulfatase activity was first assessed by measuring the release of 4-nitrophenol from potassium 4-nitrophenyl sulfate (pNP-sulfate). Briefly, reactions mixtures contained 0.75mg/ml pNP-sulfate and 1 µM enzyme in a final volume of 100µl, and were incubated at 25°C. Release of 4-nitrophenol over time was quantified at 405 nm. Screening of activity on oligo and polysaccharides was evaluated and monitored as described above. No enzyme activity was observed in heat-inactivated or enzyme-free control reactions, confirming that the observed activity was enzyme-dependent.

### Sequential degradation of chondroitin sulfate by the four enzymes of the PUL_GAG_

A reaction mix was prepared containing 0.2% CS in Tris HCl 100 mM pH7.5, NaCl 50 mM and 5 µM of *So*S1_27. The reaction was incubated at room temperature overnight. Each enzyme was successively added every 24 hours and further incubated at room temperature. Samples were taken after each 24 hours period incubation and analyzed by HPLC-SEC as described above.

### Biochemical characterization of *So*S1_27

To further characterize the activity of *So*S1_27, CS derived disaccharides with specific sulfation motifs were tested: monosulfated disaccharides at position 4 or 6 (CS4S and CS6S, respectively), disulfated disaccharides at positions 2 and 4 or 2 and 6 (CS2S4S and CS2S6S, respectively), and the trisulfated disaccharide at positions 2, 4 and 6 (CS2S4S6S). Briefly, 50 nM of purified *So*S1_27 was incubated with 50 nM of each disaccharide substrates in 50 mM Tris-HCl buffer, pH 7.5, at 37 °C, and reactions were carried out over a 0-300 min time course to monitor product formation. At each time point, the reactions were stopped by heating at 100 °C for 10 minutes, followed by centrifugation at 14000 rpm for 10 mins to remove denatured protein. The supernatants were recovered and analyzed by reverse-phase ion-pairing high-performance liquid chromatography (RPIP-HPLC) (previously described in ^44^). The desulfated products, CS0S from CS4S and CS6S from CS4S6S, were identified by comparison to standards and quantified using calibration curves. Initial rate measurements were extracted from the linear phase of product accumulation, determined from plots of product concentration versus time.

For kinetic measurements, the enzyme was incubated with varying concentrations of CS4S or CS4S6S (0–3000 µM) under the same buffer conditions, and the reaction was stopped after 9 min for CS4S, and 40 min for CS4S6S. Initial velocities (V) were calculated from the rate of product formation per minute. A Lineweaver–Burk plot (1/V=f(1/S)) was generated, and kinetic parameters Km and Vmax were estimated by linear regression of the equation 1/V=1/Vmax+(K_M_/Vmax)1/S. All assays were performed in triplicate and represent mean values ± standard deviation.

### Purification of desulfated chondroitin and ^1^H NMR analysis

First, 50 mg of commercial chondroitin sulfate were dissolved in 100 mM Tris pH 7.5, 50 mM NaCl at 0.4% and desulfated with 20 µL of SoS1_27 at 3.2 µM, for 20 h at 25 °C with shaking. After enzyme inactivation by boiling and sample filtration on 0.2 µm, the desulfated chondroitin was desalted and freeze-dried. Samples were exchanged once with D2O and were transferred into a 5 mm NMR tube. ^1^H NMR spectra were recorded at 323 K using an Advance III 400 MHz spectrometer (Bruker) operating at a frequency of 400.13 MHz for 1H. The residual signal of the solvent was used as the internal standard: HOD at 4.55 ppm. No suppression of the HOD signal was performed.

### Crystallization and structure determination of *So*S1_27

Pure and homogeneous *So*S1_27 sample was concentrated using an Amicon centrifuge filter to a final concentration of 19 mg/mL. A primary crystallization screen was carried out at the High Throughput Crystallization Laboratory (HTX Lab) of the EMBL Grenoble ^45^ using a commercial sparse-matrix screen in 96-well sitting-drop format at 20 °C. Crystals were prepared for x-ray diffraction experiments using the CrytalDirect technology ^46^. Initial crystals of *So*S1_27 were obtained in a condition containing 0.2 M magnesium formate dihydrate and 20% (w/v) PEG 3,350. Larger crystals were then produced in 24-well hanging-drop plates using the same reservoir composition, and vapor diffusion at 20 °C by mixing 1.5 µL of protein solution at 9.5 mg/mL with 1.5 µL of reservoir solution. Crystals were incubated with 2 mM CS-DP4 prepared as described in ^47^ (Supplementary Table S2). Crystals were harvested and cryo-protected by brief immersion in reservoir solution supplemented with 20% (v/v) glycerol, followed by rapid freezing in liquid nitrogen.

X-ray diffraction data were collected at the ID30A-1 beamline from the European Synchrotron Radiation Facility (ESRF) (Grenoble, France) equipped with a Pilatus (Dectris) detector ^48,49^. Datasets were indexed, integrated and scaled using the XDS package ^50^. *So*S1_27 structure was determined by molecular replacement using the program Phaser ^51^ with an Alphafold-generated model ^52^ as search coordinates. The structure refinement was performed using Phenix.refine and WinCoot ^53,54^. The final structure was validated using the servers MolProbity and CheckMyMetal ^33,55^. Figures were prepared in ChimeraX ^56^.

## Abbreviations

DP: Degree of Polymerization
DS: Dermatan Sulfate
CS: Chondroitin sulfate
HA: Hyaluronic acid
pNP: para-nitrophenol
CAZyme: Carbohydrate Active enzyme
PUL: Polysaccharide Utilization Locus
GAG: Glycosaminoglycan
GH: Glycoside Hydrolase
PL: Polysaccharide Lyase

## Acknowledgements

This work used the platforms (HTX lab) of the Grenoble Instruct-ERIC center (ISBG; UMS 3518 CNRS-CEA-UJF-EMBL) within the Grenoble Partnership for Structural Biology (PSB). This work was supported by the French National Research Agency (ANR) through a PRC scientific grant (ANR-22-CE15-0041-01), the Labex ARCANE and GRAL of the University Grenoble Alpes graduate school (Ecoles Universitaires de Recherche) CBH-EUR-GS (ANR-17-EURE-0003), FRISBI (ANR-10-INBS-05-02) and the “Investissements d’avenir” program Glyco@Alps (ANR-15-IDEX-02). IBS acknowledges integration into the Interdisciplinary Research Institute of Grenoble (IRIG, CEA). We would like to thank the staff of the ESRF and EMBL Grenoble for assistance and support in using beamline ID30A-1 under proposal number mx2498.

## Author contributions

R.V. and M.C. designed research. A.C., Y.A., M.F., M.L. performed biochemical characterization, S.D. performed NMR, L.B. contributed to chromatography analyses, C.L.B. contributed oligosaccharides, R.J.B and R.L.C performed structural characterization. R.J.B. R.L.C., R.W., R.V., W.H., M.C. analyzed and interpreted the data. M.C. drafted the manuscript. All authors contributed to the manuscript preparation and revision.

## Data availability statement (mandatory)

The atomic coordinates for the *So*S1_27 crystal structure have been deposited in the Protein Data Bank [https://www.rcsb.org/] with the PDB ID: 28MA.

## Additional Information

The authors declare no competing interests.

## Supplementary Figures

**Supplementary Figure S1: *So*1_27 is an endo-acting chondroitin 4-O-sulfatase.** A: Product profiles of oligosaccharides generated by degradation of 4S, 6S chondroitin sulfate (dashed line) or *So*S1_27 treated 4S, 6S chondroitin sulfate (solid line). B: ^1^H NMR analysis of 4S, 6S chondroitin sulfate treated or not with *So*S1_27.

**Supplementary Figure S2. Oligomeric state and conservation of the metal ion binding site in *So*S1_27.** A: The asymmetric unit comprises four chains, represented in cartoon and differentiated by color. B: Mass photometry reveals that *So*S1_27 behaves mostly as a monomer in solution. C: Structural superposition of S1 sulfatases highlights the amino acid residues coordinating metal ions at the active sites, with side chains shown as sticks. Included are SoS1_27; CS/DS-specific 4-O-sulfatases from *Bacteroides thetaiotaomicron* (PDB-ID 6S21)^15^ and *Vibrio* sp. (PDB-ID 6J66)^58^; a carrageenan-specific sulfatase from *Pseudoalteromonas fuliginea* (PDB-ID 6PSM)^59^; and a 6-O-sulfatase from *B*. *thetaiotaomicron* (PDB-ID 7P24)^60^. Note that in the three S1_27 sulfatases, three conserved residues (two aspartic acids and one histidine) participate in metal coordination. In contrast, in S1_19 and S1_11, the histidine is replaced by a polar uncharged amino acid, and an additional aspartate residue contributes to metal ion stabilization.

## Supplementary Tables

**Supplementary Table S1:**
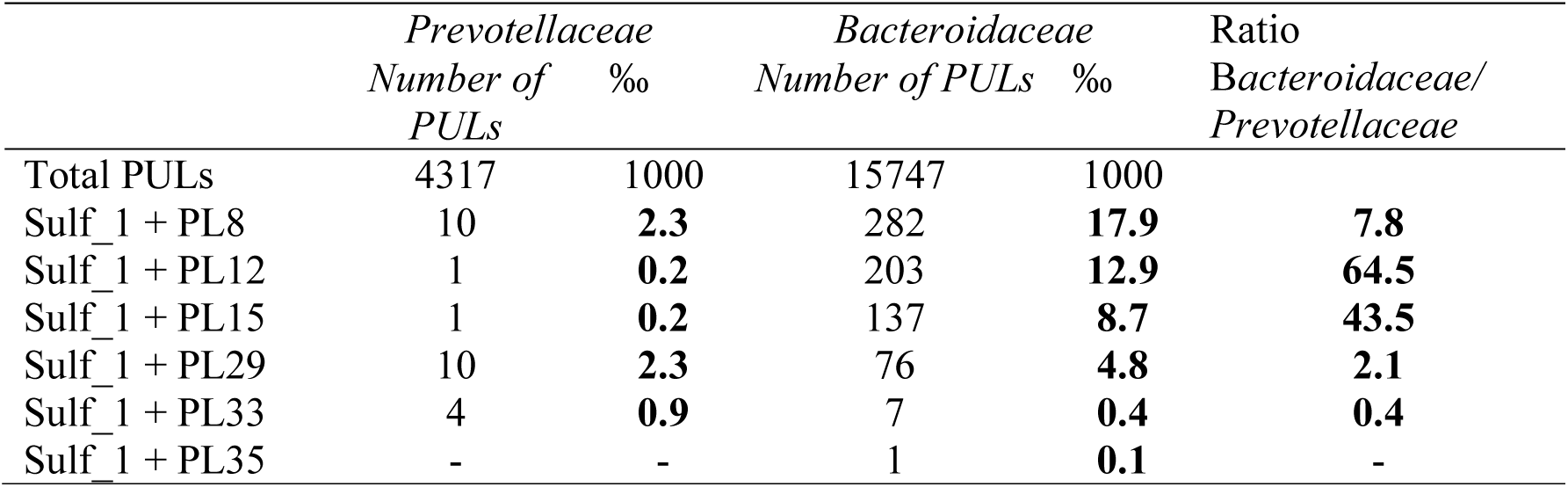
Comparative abundance of predicted GAG-targeting PUL in *Prevotellaceae* vs *Bacteroidaceae* in the PULDB.

|  | *Prevotellaceae* |  | *Bacteroidaceae* |  | Ratio |
| --- | --- | --- | --- | --- | --- |
|  | *Number of* | *‰* | *Number of PULs* | *‰* | *Bacteroidaceae/* |
|  | *PULs* |  |  |  | *Prevotellaceae* |
| Total PULs | 4317 | 1000 | 15747 | 1000 |  |
| Sulf_1 + PL8 | 10 | **2.3** | 282 | **17.9** | **7.8** |
| Sulf_1 + PL12 | 1 | **0.2** | 203 | **12.9** | **64.5** |
| Sulf_1 + PL15 | 1 | **0.2** | 137 | **8.7** | **43.5** |
| Sulf_1 + PL29 | 10 | **2.3** | 76 | **4.8** | **2.1** |
| Sulf_1 + PL33 | 4 | **0.9** | 7 | **0.4** | **0.4** |
| Sulf_1 + PL35 | - | - | 1 | **0.1** | - |

**Supplementary Table S2:** GAG poly– and oligosaccharides used in the study.

|  | Source | Osidic Composition |
| --- | --- | --- |
| **Polysaccharides (activity screening)** |  |  |
| Chondroitin sulfate | TCI | (D-GlcA β-1,3 4S D-GalNAc / 6S D-GalNAc β-1,4 )<sub>n</sub> |
| Dermatan sulfate | Glycomix | (D-IduA α-1,3 4S D-GalNAc β-1,4)<sub>n</sub> |
| Hyaluronic acid | CODIF | (D-GlcA β-1,3 D-GlcNAc β-1,4)<sub>n</sub> |
| Heparin | Carbosynth | (2S D-IduA α- /2S D-GlcA β-1,4 NS6S D-GlcN α-1,4)<sub>n</sub> |
| Heparosan | Kindly provided by B. Priem | (D-GlcA β-1,4 D-GlcNAc α-1,4)<sub>n</sub> |
| **Oligosaccharide mixes (activity screening)** |  |  |
| CS oligos | Produced in house using appropriate endo acting polysaccharide lyases | UGlcA β-1,3 4S D-GalNAc/6S D-GalNAc β-1,4 (D-GlcA β-1,3 4S D-GalNAc / 6S D-GalNAc β-1,4)<sub>n</sub> |
| HA oligos | | UGlcA β-1,3 D-GlcNAc β-1,4 (D-GlcA β-1,3 D-GlcNAc β-1,4)<sub>n</sub> |
| Heparin oligos | | UIduA α- /UGlcA β-1,4 NS6S D-GlcN α-1,4 (D-IduA α- /D-GlcA β-1,4 NS6S D-GlcN α-1,4)<sub>n</sub> |
| Heparosan oligos | | UGlcA β-1,4 D-GlcNAc α-1,4 (D-GlcA β-1,4 D-GlcNAc α-1,4)<sub>n</sub> |
| **CS disaccharides(biochemical characterization)** |  |  |
| CS4S | Iduron | UGlcA β-1,3 4S D-GalNAc |
| CS6S | | UGlcA β-1,3 6S D-GalNAc |
| CS4S6S | | UGlcA β-1,3 4S6S D-GalNAc |
| CS2S4S6S | | 2S UGlcA β-1,3 4S6S D-GalNAc |
| **CS tetrasaccharide (co-crystallization)** |  |  |
| | Synthesized as described in <sup>47</sup> | (D-GlcA β-1,3 4S6S D-GalNAc β-1,4)<sub>2</sub> |

**Supplementary Table S3:** Data collection and refinement statistics.

|  | *SoS1_27* |
| --- | --- |
| ***Data collection*** |  |
| Wavelength | 0.9655 |
| Resolution range | 60.04 - 2.39 (2.42 - 2.39) |
| Space group | P 1 |
| Unit cell | 64.49 84.85 100.91 70.85 71.9<br>86.36 |
| Total reflections | 132,074 (6,189) |
| Unique reflections | 74,555 (3,596) |
| Multiplicity | 1.8 (1.7) |
| Completeness (%) | 97.8 (95.4) |
| Mean I/sigma(I) | 2.9 (0.6) |
| Wilson B-factor | 38.65 |
| R-merge | 0.148 (1.240) |
| R-meas | 0.209 (1.751) |
| R-pim | 0.156 (0.622) |
| CC1/2 | 0.982 (0.324) |
| ***Refinement*** |  |
| Reflections used in refinement | 74188 (2482) |
| Reflections used for R-free | 3711 (116) |
| R-work | 0.2141 (0.3621) |
| R-free | 0.2746 (0.3968) |
| Number of non-hydrogen atoms | 15623 |
| macromolecules | 15576 |
| ligands | 4 |
| solvent | 43 |
| Protein residues | 1944 |
| RMS (bonds) | 0.008 |
| RMS (angles) | 1.06 |
| Ramachandran favored (%) | 96.67 |
| Ramachandran allowed (%) | 3.22 |
| Ramachandran outliers (%) | 0.10 |
| Rotamer outliers (%) | 0.96 |
| Clashscore | 4.88 |
| **Average B-factor** | 45.01 |
| *macromolecules* | 45.04 |
| *ligands* | 34.49 |
| *solvent* | 36.18 |

Statistics for the highest-resolution shell are shown in parentheses.

## References

1. Turnbaugh, P. J. et al. The human microbiome project: exploring the microbial part of ourselves in a changing world. Nature 449, 804–810 (2007).

2. Zafar, H. & Saier, M. H. Gut Bacteroides species in health and disease. Gut Microbes 13, 1–20 (2021).

3. Arumugam, M. et al. Enterotypes of the human gut microbiome. Nature 473, 174–180 (2011).

4. De Filippo, C. et al. Impact of diet in shaping gut microbiota revealed by a comparative study in children from Europe and rural Africa. Proc Natl Acad Sci U S A 107, 14691–14696 (2010).

5. Wu, G. D. et al. Linking Long-Term Dietary Patterns with Gut Microbial Enterotypes. Science 334, 105–108 (2011).

6. Dewhirst, F. E. et al. The human oral microbiome. J Bacteriol 192, 5002–5017 (2010).

7. Tuganbaev, T., Yoshida, K. & Honda, K. The effects of oral microbiota on health. Science 376, 934–936 (2022).

8. Kunath, B. J., De Rudder, C., Laczny, C. C., Letellier, E. & Wilmes, P. The oral-gut microbiome axis in health and disease. Nat Rev Microbiol 22, 791–805 (2024).

9. Montenegro, J. et al. Exploring the Influence of Gut Microbiome on Energy Metabolism in Humans. Adv Nutr 14, 840–857 (2023).

10. Tarracchini, C. et al. Vitamin biosynthesis in the gut: interplay between mammalian host and its resident microbiota. Microbiol Mol Biol Rev 89, e0018423 (2025).

11. Xu, Y., Zhu, Y., Li, X. & Sun, B. Dynamic balancing of intestinal short-chain fatty acids: The crucial role of bacterial metabolism. Trends in Food Science & Technology 100, 118–130 (2020).

12. Schütz, B. et al. Modulation of Host Immunity by Microbiome-Derived Indole-3-Propionic Acid and Other Bacterial Metabolites. Eur J Immunol 55, e202451594 (2025).

13. Yabuuchi, S. et al. Enhanced propagation of Granulicatella adiacens from human oral microbiota by hyaluronan. Sci Rep 12, 10948 (2022).

14. Rawat, P. S., Seyed Hameed, A. S., Meng, X. & Liu, W. Utilization of glycosaminoglycans by the human gut microbiota: participating bacteria and their enzymatic machineries. Gut Microbes 14, 2068367 (2022).

15. Ndeh, D. et al. Metabolism of multiple glycosaminoglycans by Bacteroides thetaiotaomicron is orchestrated by a versatile core genetic locus. Nat Commun 11, 646 (2020).

16. Fehlner-Peach, H. et al. Distinct Polysaccharide Utilization Profiles of Human Intestinal Prevotella copri Isolates. Cell Host & Microbe 26, 680–690.e5 (2019).

17. Könönen, E. & Gursoy, U. K. Oral Prevotella Species and Their Connection to Events of Clinical Relevance in Gastrointestinal and Respiratory Tracts. Front Microbiol 12, 798763 (2021).

18. Ricard-Blum, S. et al. A biological guide to glycosaminoglycans: current perspectives and pending questions. The FEBS Journal 291, 3331–3366 (2024).

19. Drula, E. et al. The carbohydrate-active enzyme database: functions and literature. Nucleic Acids Res 50, D571–D577 (2021).

20. Stam, M. et al. SulfAtlas, the sulfatase database: state of the art and new developments. Nucleic Acids Res 51, D647–D653 (2023).

21. Terrapon, N. et al. PULDB: the expanded database of Polysaccharide Utilization Loci. Nucleic Acids Res 46, D677–D683 (2018).

22. Briliūtė, J. et al. Complex N-glycan breakdown by gut Bacteroides involves an extensive enzymatic apparatus encoded by multiple co-regulated genetic loci. Nat Microbiol 4, 1571–1581 (2019).

23. Luis, A. S. et al. Dietary pectic glycans are degraded by coordinated enzyme pathways in human colonic Bacteroides. Nat Microbiol 3, 210–219 (2018).

24. Martens, E. C. et al. Recognition and degradation of plant cell wall polysaccharides by two human gut symbionts. PLoS Biol 9, e1001221 (2011).

25. Cartmell, A. et al. How members of the human gut microbiota overcome the sulfation problem posed by glycosaminoglycans. Proc Natl Acad Sci U S A 114, 7037–7042 (2017).

26. Linares-Pastén, J. A. et al. Novel xylan-degrading enzymes from polysaccharide utilizing loci of Prevotella copri DSM18205. Glycobiology 31, 1330–1349 (2021).

27. Golisch, B. et al. The molecular basis of cereal mixed-linkage β-glucan utilization by the human gut bacterium Segatella copri. J Biol Chem 300, 107625 (2024).

28. Zhou, C. et al. Glycoside hydrolase–mediated glucomannan catabolism in Segatella copri, a target of microbiota-directed foods for malnourished children. Proc Natl Acad Sci U S A 122, e2521522122.

29. Hitch, T. C. A. et al. A taxonomic note on the genus Prevotella: Description of four novel genera and emended description of the genera Hallella and Xylanibacter. Syst Appl Microbiol 45, 126354 (2022).

30. Loiodice, M. et al. Bacterial polysaccharide lyase family 33: Specificity from an evolutionarily conserved binding tunnel. Proc Natl Acad Sci U S A 122, e2421623122 (2025).

31. Krissinel, E. & Henrick, K. Inference of Macromolecular Assemblies from Crystalline State. Journal of Molecular Biology 372, 774–797 (2007).

32. Hanson, S. R., Best, M. D. & Wong, C.-H. Sulfatases: structure, mechanism, biological activity, inhibition, and synthetic utility. Angew Chem Int Ed Engl 43, 5736–5763 (2004).

33. Zheng, H. et al. CheckMyMetal: a macromolecular metal-binding validation tool. Acta Crystallogr D Struct Biol 73, 223–233 (2017).

34. Chevenier, A. et al. Structure, function and catalytic mechanism of the carrageenan-sulfatases from the marine bacterium *Zobellia galactanivorans* DsijT. Carbohydrate Polymers 358, 123487 (2025).

35. Gupta, V. K., Chaudhari, N. M., Iskepalli, S. & Dutta, C. Divergences in gene repertoire among the reference Prevotella genomes derived from distinct body sites of human. BMC Genomics 16, 153 (2015).

36. Fletcher, J. R., Malik, A., Driggers, J. & Hunter, R. C. Cross-feeding interactions between Fusobacterium nucleatum and the glycan forager Segatella oris. mSystems 11, e00922-25.

37. Accetto, T. & Avguštin, G. Polysaccharide utilization locus and CAZYme genome repertoires reveal diverse ecological adaptation of Prevotella species. Systematic and Applied Microbiology 38, 453–461 (2015).

38. Liu, G. et al. Identification and characterization of a chondroitinase ABC with a novel carbohydrate-binding module. Int J Biol Macromol 271, 132518 (2024).

39. El Masri, R., Crétinon, Y., Gout, E. & Vivès, R. R. HS and Inflammation: A Potential Playground for the Sulfs? Front Immunol 11, 570 (2020).

40. Ulmer, J. E. et al. Characterization of glycosaminoglycan (GAG) sulfatases from the human gut symbiont Bacteroides thetaiotaomicron reveals the first GAG-specific bacterial endosulfatase. J Biol Chem 289, 24289–24303 (2014).

41. Kumar, S. et al. MEGA12: Molecular Evolutionary Genetic Analysis Version 12 for Adaptive and Green Computing. Mol Biol Evol 41, msae263 (2024).

42. Edgar, R. C. MUSCLE: multiple sequence alignment with high accuracy and high throughput. Nucleic Acids Res 32, 1792–1797 (2004).

43. Couturier, M. et al. Functional exploration of the glycoside hydrolase family GH113. PLoS One 17, e0267509 (2022).

44. Laguri, C., Sadir, R., Gout, E., Vivès, R. R. & Lortat-Jacob, H. Preparation and Characterization of Heparan Sulfate-Derived Oligosaccharides to Investigate Protein-GAG Interaction and HS Biosynthesis Enzyme Activity. Methods Mol Biol 2303, 121–137 (2022).

45. Dimasi, N., Flot, D., Dupeux, F. & Márquez, J. A. Expression, crystallization and X-ray data collection from microcrystals of the extracellular domain of the human inhibitory receptor expressed on myeloid cells IREM-1. Acta Cryst F 63, 204–208 (2007).

46. Zander, U. et al. Automated harvesting and processing of protein crystals through laser photoablation. Acta Cryst D 72, 454–466 (2016).

47. Jacquinet, J.-C., Lopin-Bon, C. & Vibert, A. From polymer to size-defined oligomers: a highly divergent and stereocontrolled construction of chondroitin sulfate A, C, D, E, K, L, and M oligomers from a single precursor: part 2. Chemistry 15, 9579–9595 (2009).

48. Svensson, O., Malbet-Monaco, S., Popov, A., Nurizzo, D. & Bowler, M. W. Fully automatic characterization and data collection from crystals of biological macromolecules. Acta Crystallogr D Biol Crystallogr 71, 1757–1767 (2015).

49. Fogh, R. H. et al. Automated workflows for strategy computation and data collection at synchrotron beamlines. Acta Crystallogr D Struct Biol 82, 314–329 (2026).

50. Kabsch, W. Integration, scaling, space-group assignment and post-refinement. Acta Crystallogr D Biol Crystallogr 66, 133–144 (2010).

51. McCoy, A. J. et al. Phaser crystallographic software. J Appl Crystallogr 40, 658–674 (2007).

52. Abramson, J. et al. Accurate structure prediction of biomolecular interactions with AlphaFold 3. Nature 630, 493–500 (2024).

53. Afonine, P. V. et al. Real-space refinement in PHENIX for cryo-EM and crystallography. Acta Crystallogr D Struct Biol 74, 531–544 (2018).

54. Emsley, P., Lohkamp, B., Scott, W. G. & Cowtan, K. Features and development of Coot. Acta Crystallogr D Biol Crystallogr 66, 486–501 (2010).

55. Williams, C. J. et al. MolProbity: More and better reference data for improved all-atom structure validation. Protein Sci 27, 293–315 (2018).

56. Meng, E. C. et al. UCSF ChimeraX: Tools for structure building and analysis. Protein Sci 32, e4792 (2023).

57. Käll, L., Krogh, A. & Sonnhammer, E. L. L. Advantages of combined transmembrane topology and signal peptide prediction--the Phobius web server. Nucleic Acids Res 35, W429–432 (2007).

58. Wang, S. et al. Comparative Study of Two Chondroitin Sulfate/Dermatan Sulfate 4-O-Sulfatases With High Identity. Front. Microbiol. 10, (2019).

59. Hettle, A. G. et al. Insights into the κ/ι-carrageenan metabolism pathway of some marine Pseudoalteromonas species. Commun Biol 2, 474 (2019).

60. Luis, A. S. et al. Sulfated glycan recognition by carbohydrate sulfatases of the human gut microbiota. Nat Chem Biol 18, 841–849 (2022).

